# Genome Warehouse: A Public Repository Housing Genome-scale Data

**DOI:** 10.1101/2021.02.10.430367

**Authors:** Meili Chen, Yingke Ma, Song Wu, Xinchang Zheng, Hongen Kang, Jian Sang, Xingjian Xu, Lili Hao, Zhaohua Li, Zheng Gong, Jingfa Xiao, Zhang Zhang, Wenming Zhao, Yiming Bao

**Author notes:** Equal contribution (Meili Chen, Yingke Ma). Corresponding author. (Bao Y). Division of Cancer Epidemiology and Genetics, National Cancer Institute, National Institutes of Health, Bethesda, Maryland 20892, USA. College of Computer Science Technology, Inner Mongolia Normal University, Hohhot, Inner Mongolia 010010, China (Xingjian Xu).

## Abstract

The Genome Warehouse (GWH) is a public repository housing genome assembly data for a wide range of species and delivering a series of web services for genome data submission, storage, release, and sharing. As one of the core resources in the National Genomics Data Center (NGDC), part of the China National Center for Bioinformation (CNCB, https://bigd.big.ac.cn/), GWH accepts both full genome and partial genome (chloroplast, mitochondrion, and plasmid) sequences with different assembly levels, as well as an update of existing genome assemblies. For each assembly, GWH collects detailed genome-related metadata including biological project and sample, and genome assembly information, in addition to genome sequence and annotation. To archive high-quality genome sequences and annotations, GWH is equipped with a uniform and standardized procedure for quality control. Besides basic browse and search functionalities, all released genome sequences and annotations can be visualized with JBrowse. By December 2020, GWH has received 17,264 direct submissions covering a diversity of 949 species, and has released 3370 of them. Collectively, GWH serves as an important resource for genome-scale data management and provides free and publicly accessible data to support research activities throughout the world. GWH is publicly accessible at https://bigd.big.ac.cn/gwh/.

## Introduction

Genome sequences and annotations are fundamental information for a wide range of genome-related studies, including various omics data analysis such as genome [1], transcriptome [2], epigenome [3,4], and genome variation [5,6]. China, as one of the most biodiverse countries in the world, harbors more than 10% of the world’s known species [7]. In the past decades, a large number of genome assemblies of featured and important animals and crops in China have been sequenced [1, 8–11], most of which were submitted to International Nucleotide Sequence Database Collaboration (INSDC) members (National Center for Biotechnology Information (NCBI), European Bioinformatics Institute (EBI), and DNA Data Bank of Japan (DDBJ)) [12]. With the rapid growth of genome assembly data, in China for example, large genome data size, slow data transfer rate due to limited international network transfer bandwidth, and language barrier for communication of technical issues have obstructed researchers from efficiently submitting their data to INSDC members. All these call for a centralized genomic data repository within China to complement the INSDC.

Here, we report the Genome Warehouse (GWH, https://bigd.big.ac.cn/gwh/), a centralized resource housing genome assembly data and delivering a series of genome data services. As one of the core resources in the National Genomics Data Center (NGDC), part of the China National Center for Bioinformation (CNCB, https://bigd.big.ac.cn/) [13], the aim of GWH is to accept data submissions worldwide and provide an important resource for genome data quality control, data archive, rapid release, and public sharing (*e.g*., with INSDC) in support of research activities from all over the world. To date, GWH has received a total of 12,366 genome submissions (including 14 international submissions), demonstrating its increasingly important role in global genome data management and sharing.

## Data model

Designed for compatibility with the INSDC data model, each genome assembly in GWH is linked to a BioProject (https://bigd.big.ac.cn/bioproject) and a BioSample (https://bigd.big.ac.cn/biosample), which are two fundamental resources for metadata description in CNCB-NGDC. Full or partial (chloroplast, mitochondrion, and plasmid) genome assemblies with different assembly levels (complete, draft in chromosome, scaffold, and contig) are all acceptable and existing genome assemblies are allowed to be updated. Accession numbers are assigned with the following rules (**Figure 1**): (1) each genome assembly has an accession number prefixed with “GWH”, followed by four capital letters and eight zeros (*e.g*., GWHAAAA00000000); (2) genome sequences have the same accession number format as their corresponding genome assembly, with the exception that the eight digits start from 00000001 and increase in order (*e.g*., GWHAAAA00000001); (3) genes have similar accession pattern as those of genome sequences, with the addition of letter “G” between the GWH prefix and the four capital letters, and there are six digits at the end instead of eight (*e.g*., GWHGAAAA000001); (4) transcripts use the letter “T” to replace “G” in accession numbers for genes (*e.g*., GWHTAAAA000001); (5) proteins use the letter “P” to replace “G” in accession numbers for genes (*e.g*., GWHPAAAA000001); (6) if the submission is an update of existing submission in GWH, it will be assigned a dot and an incremental number to represent the version (*e.g*., GWHAAAA00000000.1).

**Figure 1.**
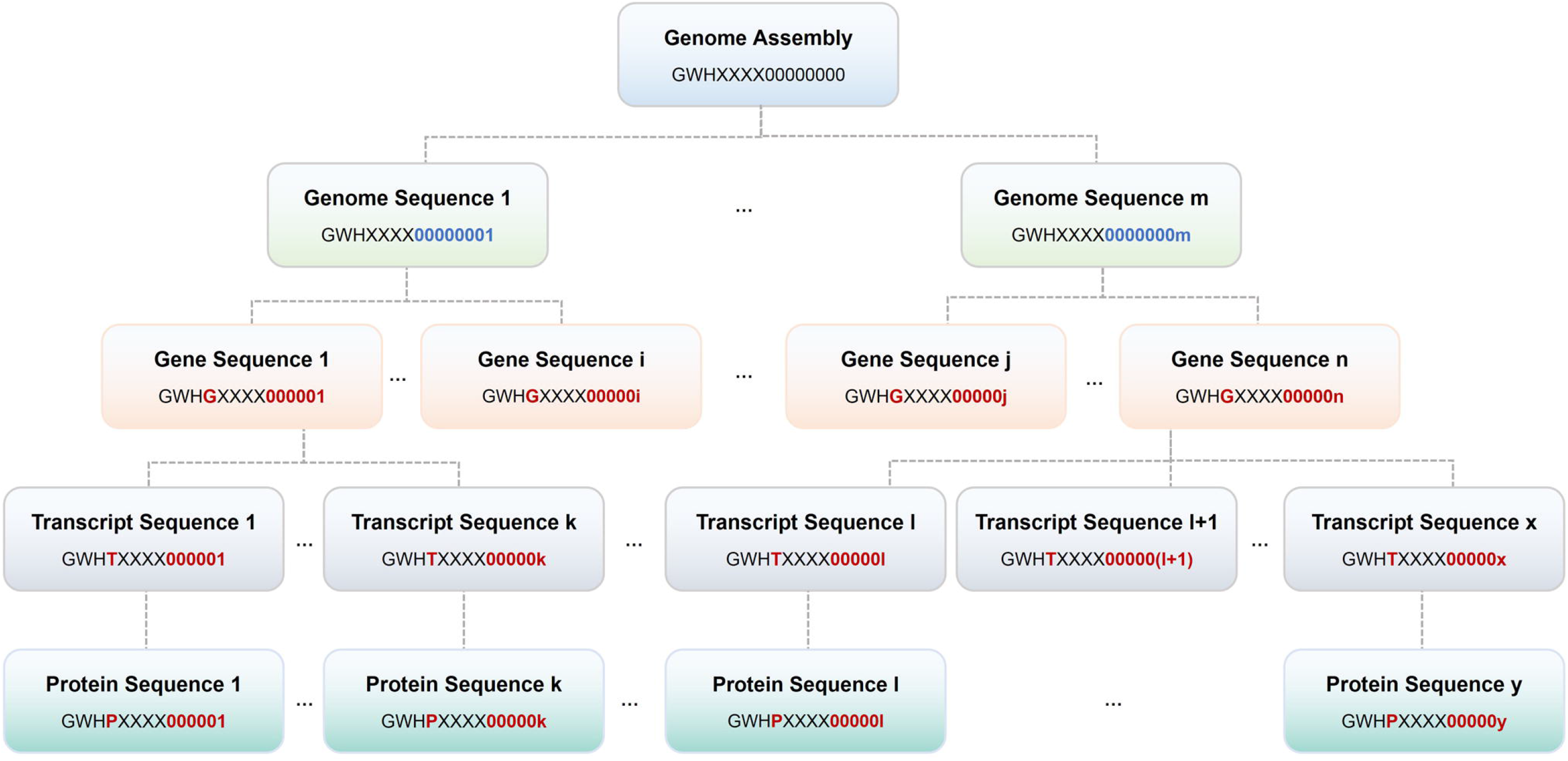
Data model in GWH. Genome assembly accession number is prefixed with “GWH”, followed by four capital letters (represented by XXXX) and 8 zeros. For genome sequence accessions, eight digits increase in order. For gene sequence, transcript sequence, and protein sequence accessions, G, T, and P are followed by the GWH prefix, respectively, with six digits at the end that increase in order.

## Database components

GWH is a centralized resource housing genome-scale data, with the purpose to archive high-quality genome sequences and annotation information. GWH is equipped with a series of web services for genome data submission, release, and sharing, accordingly involving three major components, namely, data submission, quality control, and archive and release (Figure 2).

**Figure 2.**
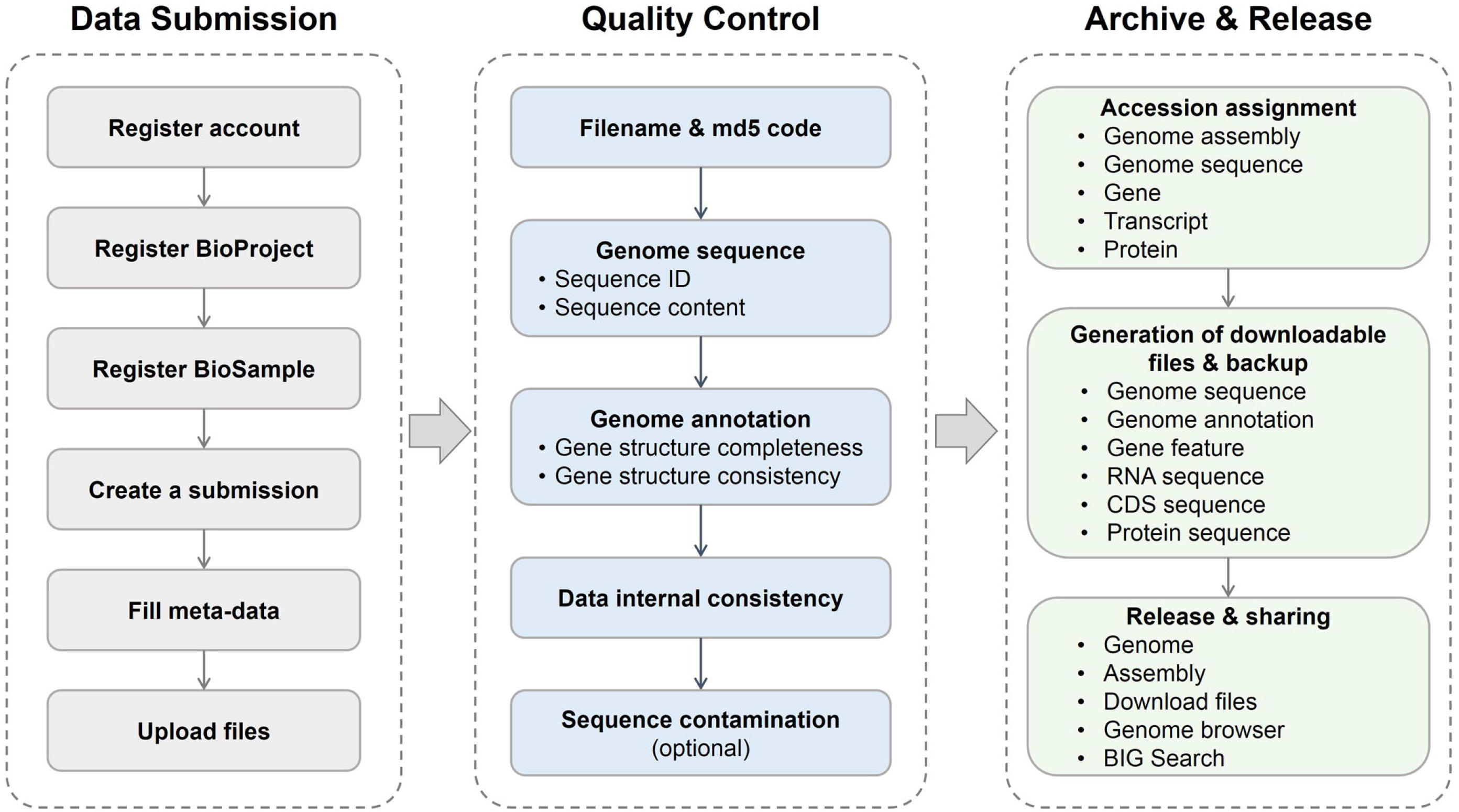
Major components in GWH data processing workflow.

### Data submission

GWH not only accepts genome assembly associated data through an on-line submission system but also allows off-line batch submissions. Users need to register first and then to provide complete description on submitted genome sequences. Biological project and sample information should be provided (through BioProject and BioSample, respectively) together with genome assembly sequence, annotation, and associated metadata. Metadata mainly consist of a variety of information about submitter, general assembly, file(s), sequence assignment, and publication (if available). After submission, GWH runs an automated quality control pipeline to check the validity and consistency of submitted genome sequence and genome annotation files. Accession numbers are assigned to assemblies and sequences upon the pass of quality control. The updated assembly data can also be submitted to GWH. It should be noted that compatible with the INSDC members (*e.g*., NCBI GenBank), it is the responsibility of the submitters to ensure the data quality, completeness, and consistency and GWH does not warrant or assume any legal liability or responsibility for the data accuracy.

### Quality control

After metadata and file(s) are received, GWH automatically runs standardized quality control (QC) to check 45 different types of errors in submitted genome sequences and annotations, and to scan for contaminated genome sequences (see details at https://bigd.big.ac.cn/gwh/documents) if needed (Figure 2), which roughly falls into 5 QC steps: (1) The component will check the consistency of file(s) according to filename and md5 code. (2) For genome sequences, the component will check the legality of genome sequence ID and sequence content, *e.g*., unique sequence ID, sequence composition (A/T/C/G or degenerate base), sequence length (≥ 200 bp). (3) For genome annotations, the component will check gene structure completeness and consistency, *e.g*., unique ID, a exon/CDS/UTR coordinate falling within the corresponding gene coordinate, strand consistency for all features (including gene/transcript/exon/CDS/UTR), codon validity (*e.g*., valid start/stop codon, no internal stop codon). (4) Finally, it will check the internal consistency of genome sequence and annotation, *e.g*., sequence ID in genome annotation must match genome sequence ID, a feature coordinate falling within the range of the corresponding genome sequence. (5) Genome sequences will also be scanned to check vectors, adaptors, primers, and indices (collected from UniVec database, ftp://ftp.ncbi.nlm.nih.gov/pub/UniVec/) using NCBI’s VecScreen (https://www.ncbi.nlm.nih.gov/tools/vecscreen/). If there is an error, a report will be automatically sent to the submitter by email. To finish a successful submission, the submitter needs to fix all errors and resubmit files until they pass the QC process.

### Archive and release

GWH will assign a unique accession number to the submitted genome assembly upon the pass of quality control, allot accession numbers for each genome sequence, gene, transcript, and protein, generate and backup downloadable files of genome sequence and annotation in FASTA, GFF3, and TSV formats. Data generation is performed with in-house-writing scripts based on submitted genome sequence and annotation files. In order to ensure the security of submitted data, a copy of backup data is stored on a physically separate disk. GWH will release sequence data on a user-specified date, unless a paper citing the sequence or accession number is published prior to the specified release date, in which case the sequence will be released immediately. For the released data, GWH will generate web pages containing two primary tables: genome and assembly. The former shows species taxonomy information and genome assemblies, and the latter contains general information of the assembly (including external links to other related resources), statistics of genome assembly and its corresponding annotation. All released data are publicly available at GWH FTP site (ftp://download.big.ac.cn/gwh/). GWH provides data visualization for both genome sequence and genome annotation using JBrowse [14]. It offers statistics and charts in light of total holdings, assembly levels, genome representations, citing articles, submitting organizations, sequencing platforms, assembly methods, and downloads. GWH provides user-friendly web interfaces for data browse and query using BIG Search [13], in order to help users find any released data of interest. For a released genome assembly, GWH also provides machine-readable APIs (Application Programming Interfaces) for publicly sharing and automatically obtaining information on its associated BioProject, BioSample, genome, and assembly metadata and file paths.

### Global sharing of SARS-CoV-2 and coronavirus genomes

During the COVID-19 outbreak, GWH, in support of the 2019 Novel Coronavirus Resource (2019nCoVR) [15, 16] has received worldwide submissions of more than a thousand SARS-CoV-2 genome assemblies with standardized genome annotations [17], and has released 134 of them. To expand the international influence of data, 62 of the released sequences have been shared, with the submitters’ permission, in GenBank [18] through a data exchange mechanism established with NCBI. In this model, GWH accessions are represented as secondary accessions in NCBI GenBank records, which are retrievable by the NCBI Entrez system. This model sets a good example for data sharing among different data centers.

In addition, GWH offers sequences of the Coronaviridae family to facilitate researchers to reach the data conveniently and thus to study the relationship between SARS-CoV-2 and other coronaviruses. To promote the data sharing and make all relevant information of the Coronaviridae readily available, GWH integrates genomic and proteomic sequences as well as their metadata information from NCBI [19], China National GeneBank Database (CNGBdb) [20], National Microbiology Data Center (NMDC) [21] and CNCB-NGDC. Duplicated records from different sources are identified and removed to gain a non-redundant dataset. As of December 31, 2020, the dataset has 83,095 nucleotide and 575,438 protein sequences of the Coronaviridae. Filters are implemented to narrow down the required Coronaviridae sequences using multiple conditions, including country/region, host, isolation source, length, and collection date. Both the metadata and sequences of the filtered results can be selected and downloaded as a separate file. The daily updated sequences and all sequences can also be downloaded from FTP (ftp://download.big.ac.cn/Genome/Viruses/Coronaviridae/).

### Data statistics

By December, 2020, GWH has received 17,264 direct submissions covering a broad diversity of species (**Table 1**) with different assembly levels (Figure 3). These genome assemblies link to 301 BioProjects and 16,538 BioSamples, and are submitted by 231 submitters from 61 institutions (including 5 international submitters from 2 countries). There are a total of 3370 released submissions, which were reported in 83 articles from 44 journals. GWH has over 135,000 visits from 153 countries/regions, with ∼891,000 downloads. The amount of data, visits, and downloads in the GWH has been on the dramatic increase over the past years, clearly showing its great utility in genome-scale data management.

**Table 1.**
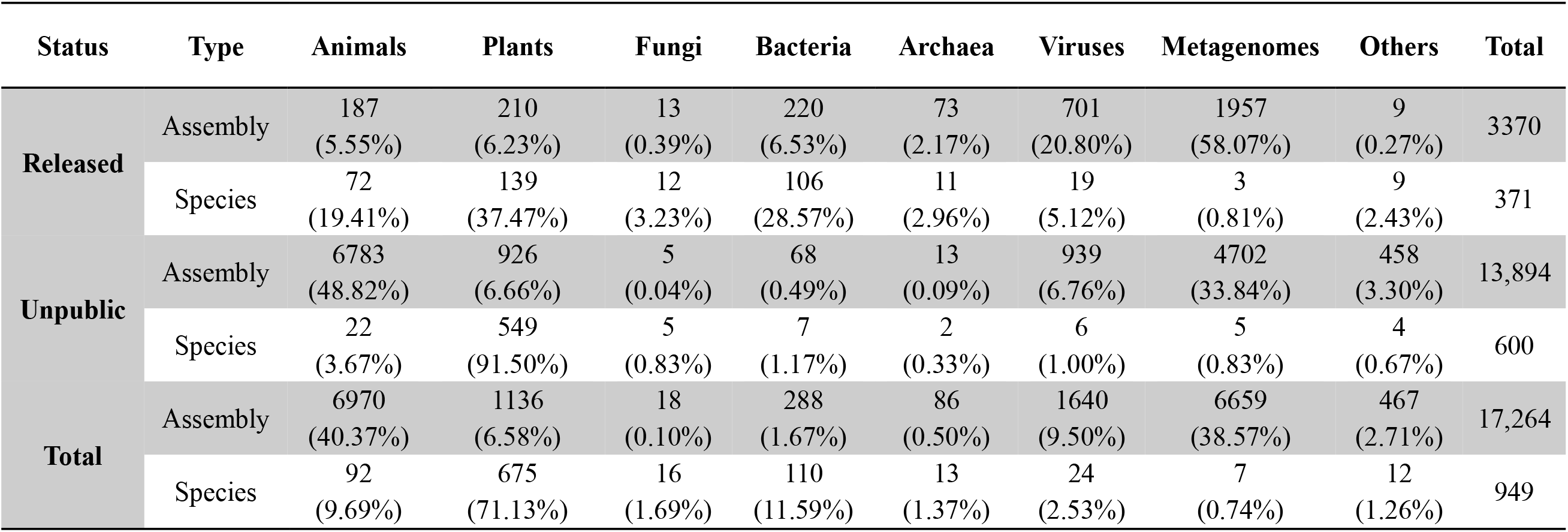
Total data holdings in GWH.

**Figure 3.**
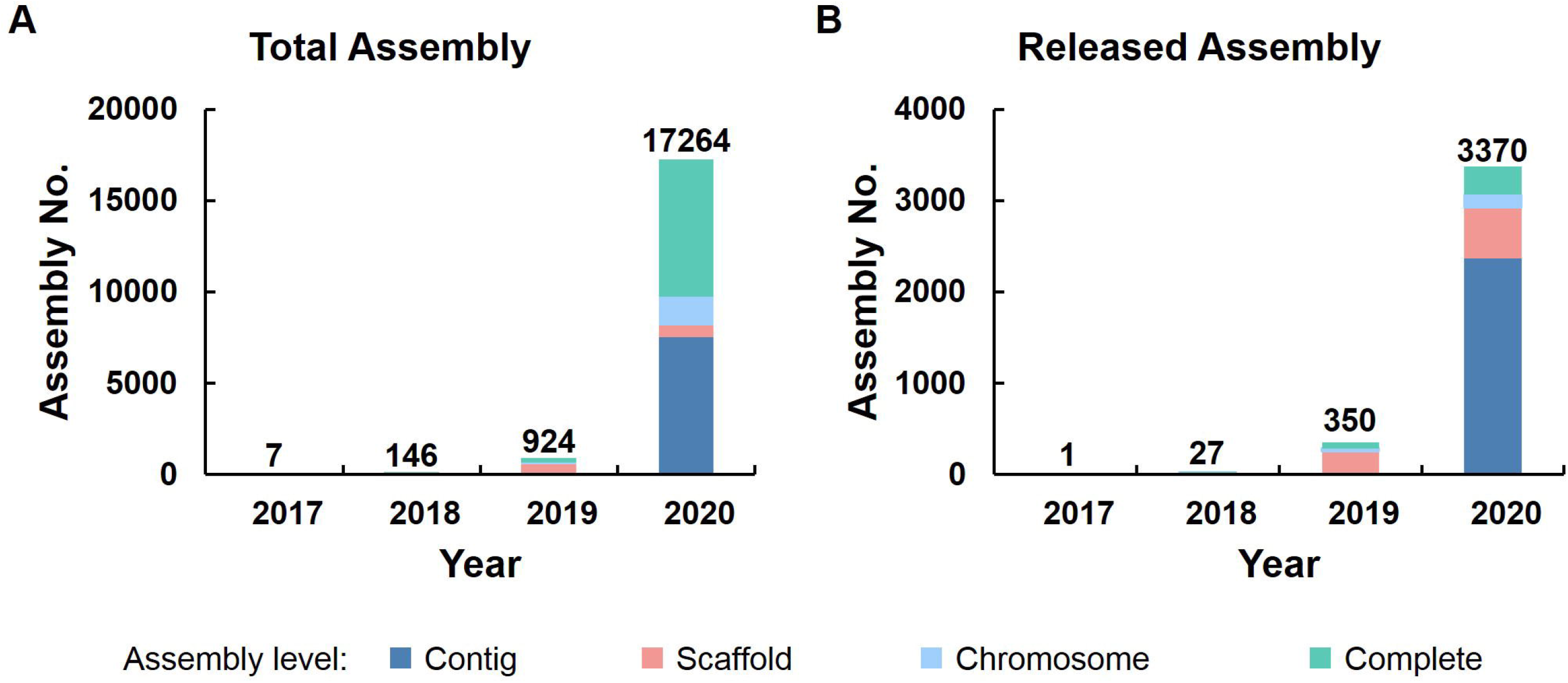
Statistics of genome assembly in GWH (as of December 31, 2020)

### Summary and future directions

Collectively, GWH is a user-friendly portal for genome data submission, release, and sharing associated with a matched series of services. The rapid growth of genome assembly submissions demonstrates the great potential of GWH as an important resource for accelerating the worldwide genomic research. With the aim to fully realize the findability, accessibility, interoperability, and reusability (FAIR) of genome data [22], GWH has made ongoing efforts, including but not limited to, improvement of web interfaces for data submission, presentation, and visualization, continuous integration of newly sequenced genomes, and development of useful online tools to help users analyse genome data (such as BLAST [23]). Therefore, we will put in more efforts to provide genome annotation services, especially for bacteria and archaea genomes, with the particular consideration that uniform standardized annotation determines the accuracy of downstream data analysis. Besides, we will expand the Coronaviridae dataset to other important pathogens to improve the ability of public health emergency response. Finally, we plan to share and exchange all public genome assembly data with the INSDC members to provide comprehensive data for researchers globally.

## CRediT author statement

**Meili Chen:** Methodology, Software, Investigation, Data Curation, Writing - Original Draft, Project administration. **Yingke Ma:** Software, Writing - Original Draft. **Song Wu:** Software, Data Curation. **Xinchang Zheng:** Data Curation. **Hongen Kang:** Software. **Jian Sang:** Investigation, Data Curation. **Xingjian Xu:** Software. **Lili Hao:** Investigation. **Zhaohua Li:** Data Curation. **Zheng Gong:** Data Curation. **Jingfa Xiao:** Writing - Review & Editing. **Zhang Zhang:** Writing - Review & Editing. **Wenming Zhao:** Writing - Review & Editing. **Yiming Bao:** Conceptualization, Writing - Review & Editing, Supervision.

## Acknowledgments

We thank Profs. Jingchu Luo and Weimin Zhu for their helpful suggestions and a number of users for reporting bugs and sending comments. We also thank the NCBI GenBank group, especially Ilene Mizrachi, Karen Clark, Mark Cavanaugh, and Linda Yankie, for their valuable advices on sequence contamination scanning and SARS-CoV-2 sequence exchange. This work was supported by Strategic Priority Research Program of Chinese Academy of Sciences [XDB38060100 and XDB38030200 to YB; XDB38050300 to WZ; XDB38030400 to JX; XDA19050302 to ZZ]; National Key Research and Development Program of China [2016YFE0206600 to YB; 2020YFC0847000, 2018YFD1000505, 2017YFC1201202, and 2016YFC0901603 to WZ; 2017YFC0907502 to ZZ]; The 13th Five-year Informatization Plan of Chinese Academy of Sciences [XXH13505-05 to YB]; Genomics Data Center Construction of Chinese Academy of Sciences [XXH-13514-0202 to YB]; Open Biodiversity and Health Big Data Initiative of IUBS [to YB]; The Professional Association of the Alliance of International Science Organizations [ANSO-PA-2020-07 to YB]; National Natural Science Foundation of China [32030021 and 31871328 to ZZ]; International Partnership Program of the Chinese Academy of Sciences [153F11KYSB20160008 to ZZ].

## References

[1] Liu Y, Du H, Li P, Shen Y, Peng H, Liu S, et al. Pan-genome of wild and cultivated soybeans. Cell 2020;182:162-76.e13.

[2] Guan Y, Chen M, Ma Y, Du Z, Yuan N, Li Y, et al. Whole-genome and time-course dual RNA-Seq analyses reveal chronic pathogenicity-related gene dynamics in the ginseng rusty root rot pathogen Ilyonectria robusta. Sci Rep 2020;10:1586.

[3] Li R, Liang F, Li M, Zou D, Sun S, Zhao Y, et al. MethBank 3.0: a database of DNA methylomes across a variety of species. Nucleic Acids Res 2018;46:D288–D95.

[4] Xiong Z, Li M, Yang F, Ma Y, Sang J, Li R, et al. EWAS Data Hub: a resource of DNA methylation array data and metadata. Nucleic Acids Res 2020;48:D890–D5.

[5] Song S, Tian D, Li C, Tang B, Dong L, Xiao J, et al. Genome Variation Map: a data repository of genome variations in BIG Data Center. Nucleic Acids Res 2018;46:D944–D9.

[6] Tang B, Zhou Q, Dong L, Li W, Zhang X, Lan L, et al. iDog: an integrated resource for domestic dogs and wild canids. Nucleic Acids Res 2019;47:D793–D800.

[7] McBeath J, McBeath JH. Biodiversity conservation in China: policies and practice. Journal of International Wildlife Law & Policy 2006;9:293–317.

[8] Fan H, Wu Q, Wei F, Yang F, Ng BL, Hu Y. Chromosome-level genome assembly for giant panda provides novel insights into Carnivora chromosome evolution. Genome Biol 2019;20:267.

[9] Xia Q, Zhou Z, Lu C, Cheng D, Dai F, Li B, et al. A draft sequence for the genome of the domesticated silkworm (Bombyx mori). Science 2004;306:1937–40.

[10] Lin T, Xu X, Ruan J, Liu SZ, Wu SG, Shao XJ, et al. Genome analysis of Taraxacum kok-saghyz Rodin provides new insights into rubber biosynthesis. Natl Sci Rev 2018;5:78–87.

[11] Li C, Song W, Luo Y, Gao S, Zhang R, Shi Z, et al. The HuangZaoSi maize genome provides insights into genomic variation and improvement history of maize. Mol Plant 2019;12:402–9.

[12] Arita M, Karsch-Mizrachi I, Cochrane G. The international nucleotide sequence database collaboration. Nucleic Acids Res 2021;49:D121–D4.

[13] Members C-N, Partners. Database resources of the National Genomics Data Center, China National Center for Bioinformation in 2021. Nucleic Acids Res 2021;49:D18–D28.

[14] Buels R, Yao E, Diesh CM, Hayes RD, Munoz-Torres M, Helt G, et al. JBrowse: a dynamic web platform for genome visualization and analysis. Genome Biol 2016;17:66.

[15] Zhao WM, Song SH, Chen ML, Zou D, Ma LN, Ma YK, et al. The 2019 novel coronavirus resource. Yi Chuan 2020;42:212–21.

[16] Song S, Ma L, Zou D, Tian D, Li C, Zhu J, et al. The global landscape of SARS-CoV-2 genomes, variants, and haplotypes in 2019nCoVR. Genomics, Proteomics & Bioinformatics 2020. [DOI: https://doi.org/10.1016/j.gpb.2020.09.001]

[17] Shean RC, Makhsous N, Stoddard GD, Lin MJ, Greninger AL. VAPiD: a lightweight cross-platform viral annotation pipeline and identification tool to facilitate virus genome submissions to NCBI GenBank. BMC Bioinformatics 2019;20:48.

[18] Sayers EW, Cavanaugh M, Clark K, Ostell J, Pruitt KD, Karsch-Mizrachi I. GenBank. Nucleic Acids Res 2020;48:D84–D6.

[19] Sayers EW, Beck J, Bolton EE, Bourexis D, Brister JR, Canese K, et al. Database resources of the National Center for Biotechnology Information. Nucleic Acids Res 2021;49:D10–D7.

[20] Chen FZ, You LJ, Yang F, Wang LN, Guo XQ, Gao F, et al. CNGBdb: China National GeneBank DataBase. Yi Chuan 2020;42:799–809.

[21] Wu L, Sun Q, Desmeth P, Sugawara H, Xu Z, McCluskey K, et al. World data centre for microorganisms: an information infrastructure to explore and utilize preserved microbial strains worldwide. Nucleic Acids Res 2017;45:D611–D8.

[22] Zhang Z, Song S, Yu J, Zhao W, Xiao J, Bao Y. The elements of data sharing. Genomics Proteomics Bioinformatics 2020;18:1–4.

[23] Altschul SF, Madden TL, Schaffer AA, Zhang J, Zhang Z, Miller W, et al. Gapped BLAST and PSI-BLAST: a new generation of protein database search programs. Nucleic Acids Res 1997;25:3389–402.

